# Automated flight interception traps for interval sampling of insects

**DOI:** 10.1101/2020.02.10.941740

**Authors:** Janine Bolliger, Marco Collet, Michael Hohl, Beat Wermelinger, Martin K. Obrist

## Abstract

Recent debates on insect decline require sound assessments on the relative drivers that may negatively impact insect populations. Often, baseline data rely on insect monitorings that integrates catches over long time periods. If, however, effects of time-critical environmental factors (e.g., light pollution) are of interest, higher temporal resolution of insect observations during specific time intervals are required (e.g., between dusk and dawn). Conventional time-critical insect trapping is labour-intensive (manual activation/deactivation) and temporally inaccurate as not all traps can be serviced synchronically at different sites. Also, temporal shifts of environmental conditions (e.g., sunset/sunrise) are not accounted for.

We provide a battery-driven automated insect flight-interception trap which samples insects during seven user-defined time intervals. A commercially available flight-interception trap is fitted to a rotating platform containing seven flasks and a passage hole. The passage hole avoids unwanted sampling during time-intervals not of interest. Comparisons between two manual and two automated traps during 71 nights in 2018 showed no difference in caught insects. An experiment using 20 automated traps during 104 nights in 2019 proved that the automated flight interception traps is reliable.

The automated trap opens new research possibilities as any insect-sampling interval can be defined. It is efficient and saves manpower and associated costs as activation/deactivation is required only every seven sampling intervals. Also, the traps are accurate because all traps sample exactly the same intervals. The trap is low maintenance and robust due to straightforward technical design. It can be controlled manually or via Bluetooth and smartphone.

## Introduction

Passive traps are widely applied in documenting and monitoring insect biodiversity [1, 2]. Among traps, transparent flight-interception traps have been successfully used for many flying insect orders [3-7]. For most trap applications such as monitoring, insect sampling does not require a high temporal resolution and traps are put in place for longer time periods, typically days to weeks [6]. However, in the context of large-scale insect decline, knowledge on how individual environmental drivers act on the populations are needed [8, 9]. One of these drivers, artificial light at night and its mostly negative impacts on insects, has triggered interest among researchers in sampling during defined time intervals (i.e., nights) [10, 11]. While sampling during specific time-intervals is well established for other organisms, e.g. acoustic signals in bats [12], we are not aware of interval-sampling devices for insects. To date, two trap visits per defined time interval were required: trap activation at the beginning of the sampling interval and collection of the insects and deactivation at the end of the time interval. Apart from being labour-intensive, such sampling of time intervals is inherently inaccurate with respect to sampling synchrony because handling the traps takes time and often, experiments consist of numerous traps at different, often distant sites. In addition, seasonal time shifts of narrow time intervals such as sunset or sunrise cannot be accounted for. Lastly, given that vegetation periods are often short anyway, sample size may be even lower than desired because manpower is limited during weekends and public holidays.

We present a modified and automated insect flight interception trap whose design, construction and effectiveness allows for sampling user-defined time intervals. While the standard commercial flight interception trap remains to ensure comparable trappability (Polytrap ® https://cahurel-entomologie.com/shop/pieges/434-piege-polytrap.html), we add an automated device to extend on the storing capacity of the sampled insects. The automated sampling device (1) allows for any user-defined sampling intervals; (2) is very efficient as site visits to empty and re-activate the traps are required only after seven consecutive sampling intervals (e.g., even nights); (3) allows for accurate and synchronized exposure times of each of the seven sampling flasks in each trap; (4) is reliable and robust due to straightforward mechanical and electronical design and controllable via Bluetooth and a smartphone.

## Material, methods and results of trap evaluations

### Trap design

Our automated insect trap relies on a commercial flight interception trap (Polytrap ® https://cahurel-entomologie.com/shop/pieges/434-piege-polytrap.html) whose transparent and light plastic (PET) construction consists of two crossed PET sheets which represent an omnidirectional barrier for flying insects, a trap roof, and a funnel and beaker filled with liquid to collect the insects (Fig. 1). These traps are broadly applied in entomological research [5, 6]. While the trappability of the commercial insect trap remains the same, the actual insect sampling is automated and optimized: a platform containing seven flasks is rotated by a stepper motor which is fed by a rechargeable battery (Fig. 2a). The seven flasks contain water - if applicable - mixed with a biocide (e.g., Rocima) and allow to collect insects during seven user-defined time intervals. The time intervals are defined and stored in a table. Using two magnets and a Hall-effect sensor built into the print, the exact position of each flask can be determined. In addition to the seven flasks, the platform contains a passage hole (Fig. 2b). The hole’s position is selected during intervals where insects are not trapped. This ensures that any insect falling into the funnel during non-trapping times is immediately released via the hole (Fig. 2b; see also https://www.youtube.com/watch?v=aGFRxYjO1rM). The insect trap can be connected and controlled via Bluetooth with an App on a smart phone. For this the app BGX Commander needs to be available on the smartphone (Appendix A). Details on the electronic scheme and the state chart of the electronic software are shown in Appendices B and C.

**Figure 1.**
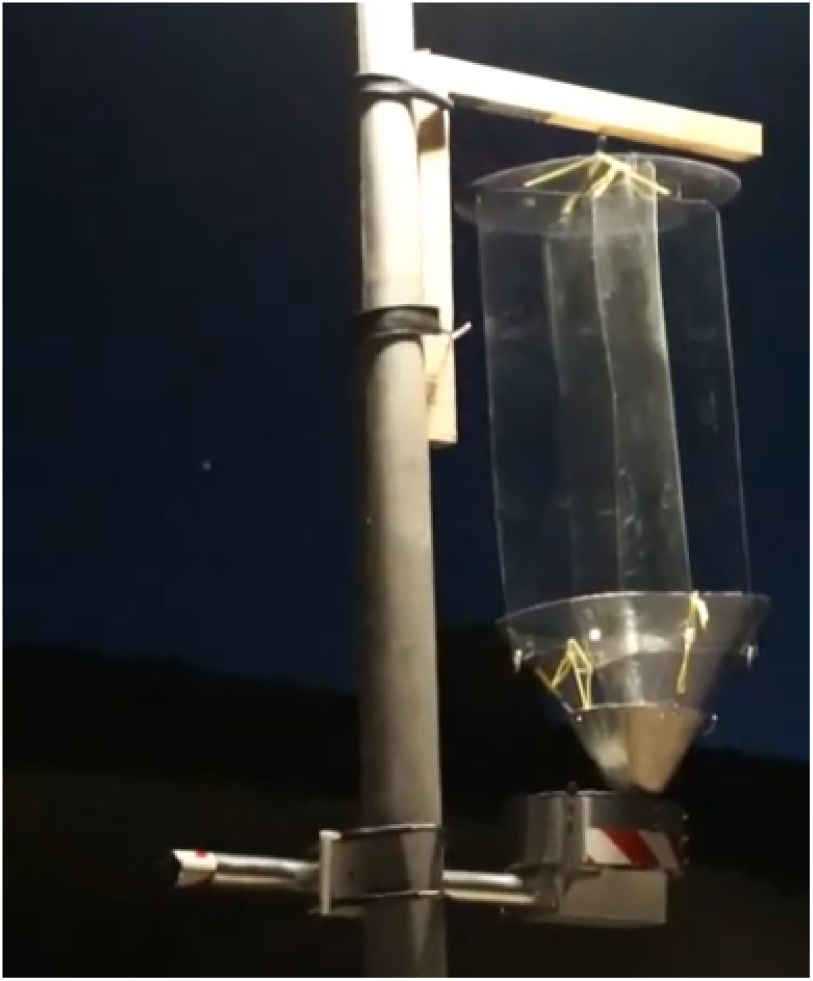
The automated insect trap mounted on a street-light pole. While the commercially available transparent flight interception trap remains unmodified, the insect sampling unit is optimized: a platform containing seven flasks is rotated by a stepper motor fed by a rechargeable battery. The seven flasks contain a liquid (water with/without biocide) and collect insects during seven user-defined time intervals (e.g., nights).

**Figure 2.**
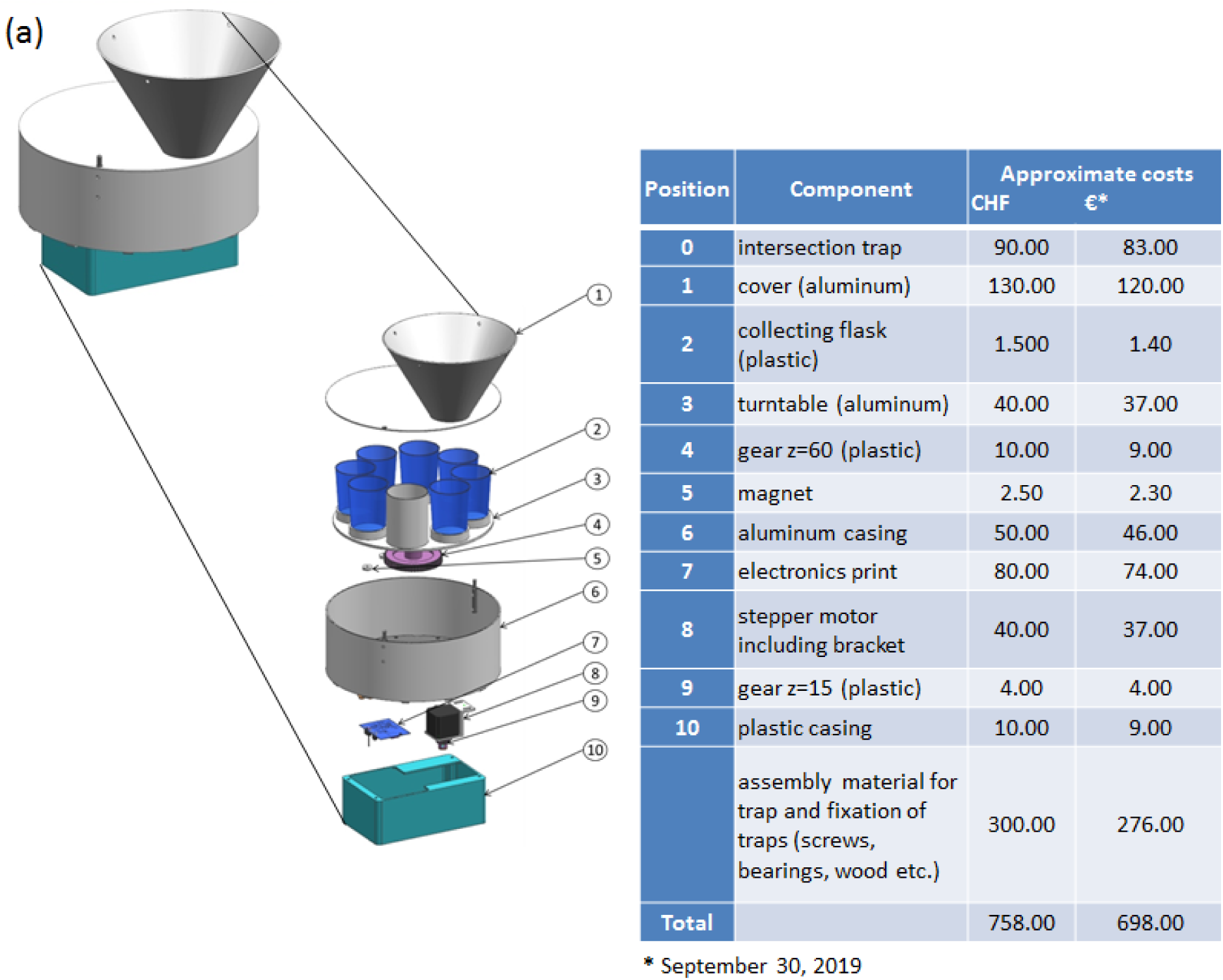
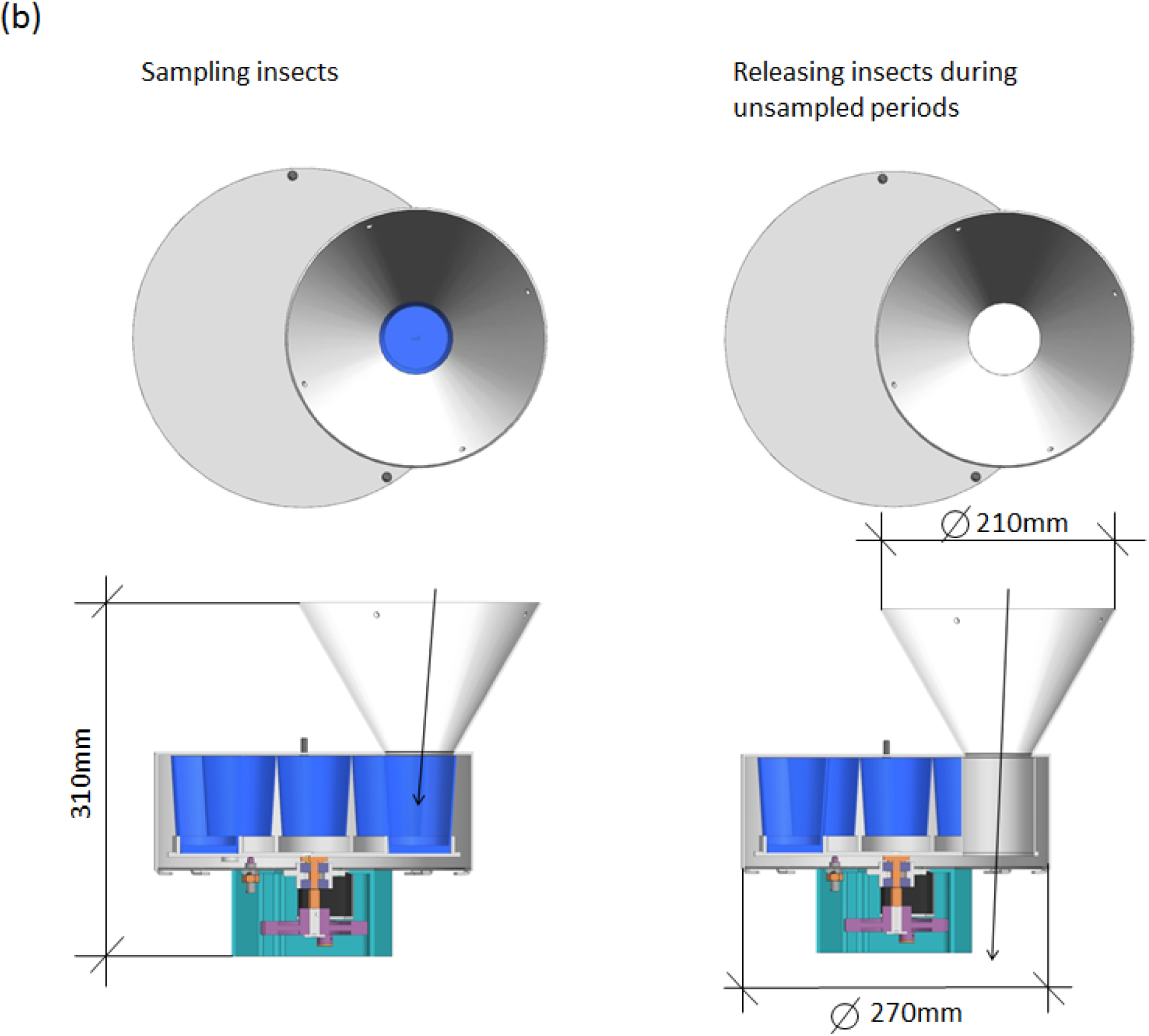
**(a)** Design and components of the platform to optimize insect sampling including a list of required components and their approximate costs. Labor costs are not included; **(b)** position during periods of sampling (flasks placed underneath funnel) and non-sampling periods, when insects are released through a passage hole.

The automated insect platform could be built larger and thus contain more flasks than just seven. Sampling seven time intervals represented a trade-off between trap and flask size, trap weight, and servicing intervals. As the traps are serviced only once a week, the flasks had to be large enough to take up enough water in order to counteract evaporation during hot weather periods.

Initial problems with humidity in the control systems of the traps was overcome by drilling ventilation holes and insulating the electronical parts against humidity (Appendix A).

### Trap evaluation

We compared the automated insect interval trap with the original manually operated trap to test the feasibility of the trap’s design and the reliability of the electronical and mechanical components. In a pilot study, insects were caught at street light poles approx. 30 m apart from each other during 71 nights between June and mid-July 2018 in the town of Birmensdorf (ZH), Switzerland, using two automated insect traps and two manually operated traps. The caught insects were sorted into six groups (Coleoptera, Diptera, Lepidoptera, Hymenoptera, Heteroptera, Neuropterida). In total, we sampled 5716 insects of which 2725 individuals were sampled using the manually operated traps and 2991 by the automated traps. There are no systematic differences between the two trap types (Fig. 3). Although comparing only two automated and two manually operated traps do not allow for formal statistical inference, the pilot study confirmed that our approach to automated insect sampling is technically feasible and reliable, resulting in comparable amounts of caught insects. A larger experiment was conducted with 20 automated insect traps during 104 nights in 2019 in the town of Weiningen (ZH), Switzerland. The traps were mounted directly to the street-light poles at a height of ca. 3 m above ground. Between May 20 and August 31 2019, we caught 49’613 insects with an overall mean of about 23 insects per night. Had we only relied on sampling of the nights during the worktime of lab assistants (only Monday-Friday instead of the full seven nights per week), we would have sampled only 60 instead of 104 nights, thus substantially increasing the insect samples by 42%. Overall, only one trap repeatedly failed and had to be replaced. All traps were restarted weekly to readjust their time measurements. Apart from humidity issues in the electronical part of the traps that could be solved (Appendix A), we did not encounter problems and the traps run smoothly for more than three months.

**Figure 3.**
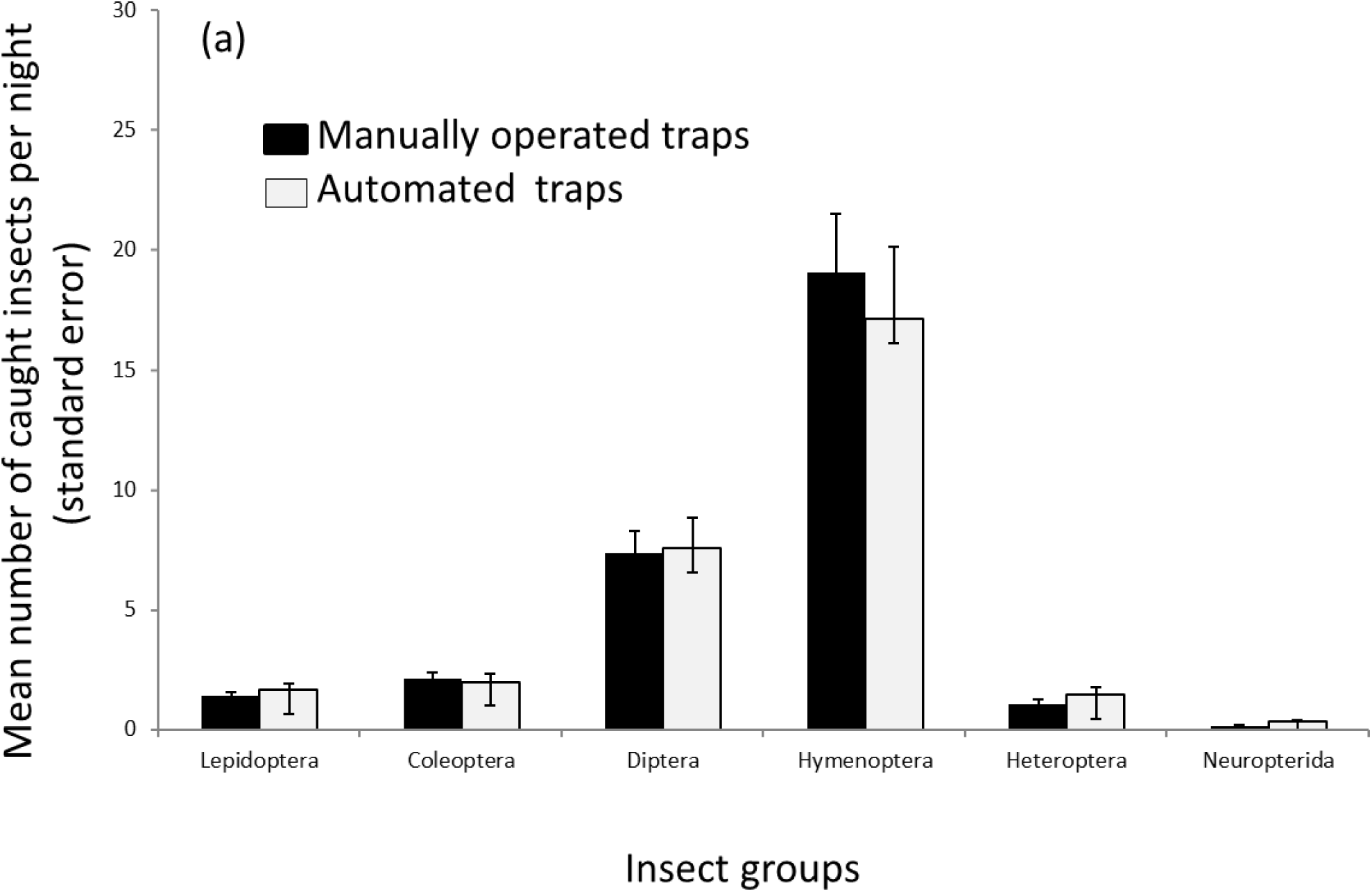
Comparison of the mean number of caught insects for automated and manually operated insect traps during 71 nights (data from summer 2018) for two traps.

## Discussion and conclusion

Insect biodiversity is threatened worldwide and not only specialist species with narrow ecological niches are affected, but also generalist species [8, 13] in protected areas [14]. To develop mitigation strategies for sustainable futures, we need more detailed knowledge on individual drivers of insect decline [8, 9]. As a step forward, we provide an automated insect flight interception trap which allows to sample insects during seven consecutive, user-defined time intervals. The time intervals may allow to identify more precisely the effects of individual environmental drivers such as e.g., street lights which likely impact nocturnal insects. The automated time-interval insect trap addresses several sampling challenges, while ensuring the same trappability as an existing and tested commercial flight interception trap: 1) the trap can be programmed to sample insects during any time interval of interest, e.g., from sunset to sunrise, including adjusted sampling-time intervals to changing day-lengths during the sampling period. This enhances sampling accuracy and saves time and personnel costs; 2) the trap connects to smartphones and can be controlled via Bluetooth (Appendix A); 3) as all traps start and terminate sampling automatically and simultaneously, the comparability between (distant) experimental sites is ensured; 4) the straightforward mechanical and electronical design of the sampling units are well accessible and easy to handle; 5) the rotating sampling platform can be mounted to any other trapping device [7].

We believe that advantages in the context of increasing environmental stressors, standardized, user-defined sampling intervals become increasingly important and supplement information obtained from long-term insect monitoring. Our automated insect trap reduces maintenance efforts and allows for a broad range of timely applications to assess environmental impacts on insect diversity. The trap is applicable to any research where insect sampling for specified time intervals is asked for. For example, we envision application to identify the natural variation of insect abundance given different treatments such as temperature, precipitation or explicit stressors in an experimental setting, e.g., application of pollutants during different development stages of insects.

The traps are with roughly 700 Euros relatively expensive. However, as the automated traps will save considerable personal and maintenance funds, the investment will pay off in the long run. Also, the price of the traps is in the same range as other electronic outdoor devices of the data-logger sector (e.g., https://www.msr.ch/en/product/msr145 for lux, temperature and relative humidity measurements).

## Author contributions

JB and MKO identified the need and developed an idea for an automated insect trap. BW provided accessibility to the entomology lab and analysed the insect data. MC and MH conceptualized and built the traps. MC and MH provided the technical figures and tables for the paper. JB, MKO and BW acquired funding for building the automated insect traps. JB provided a first draft of the paper. The authors commented on the drafts and approved the final version of this manuscript.

## Data availability

We currently do not think that it is of interest to make the insect data collected in 2018 available to the public. If, however, there is a need for this, we will gladly do so.

